# Cell-specific and shared enhancers control a high-density multi-gene locus active in mammary and salivary glands

**DOI:** 10.1101/2023.02.06.527373

**Authors:** Hye Kyung Lee, Michaela Willi, Chengyu Liu, Lothar Hennighausen

**Author notes:** Correspondence to: H.K.L and L.H.

## Abstract

Regulation of high-density loci harboring genes with different cell-specificities remains a puzzle. Here we investigate a locus that evolved through gene duplication^1^ and contains eight genes and 20 candidate regulatory elements, including a super-enhancer. Five genes are expressed in mammary glands and account for 50% of all mRNAs during lactation, two are salivary-specific and one has dual specificity. We probed the function of eight candidate enhancers through experimental mouse genetics. Deletion of the super-enhancer led to a 98% reduced expression of *Csn3* and *Fdcsp* in mammary and salivary glands, respectively, and Odam expression was abolished in both tissues. The other three casein genes were only marginally affected. Notably, super-enhancer activity requires the additional presence of a distal *Csn3*-specific enhancer. Our work identifies an evolutionary playground on which regulatory duality of a multigene locus was attained through an ancestral super-enhancer active in mammary and salivary tissue and gene-specific mammary enhancers.

## Introduction

The expansion of the secretory calcium-binding phosphoprotein (SCPP) gene family was key to the success of milk and saliva^2,3^, and its expression occurs uniquely in mineralized tissues, like mammary and salivary glands^3^. Yet, understanding enhancer structures, their specificities and interplay in complex genetic loci enabling distinct gene activities in mammary and salivary glands remains to be understood. We addressed this question experimentally in a locus harboring eight genes uniquely expressed in mammary or salivary glands or both (Fig. 1a)^1,3,4^. Duplication of *Odam*, the founding member of this family, followed by gene expansion led to a locus conserved in the mammalian lineage.

**Fig. 1.**
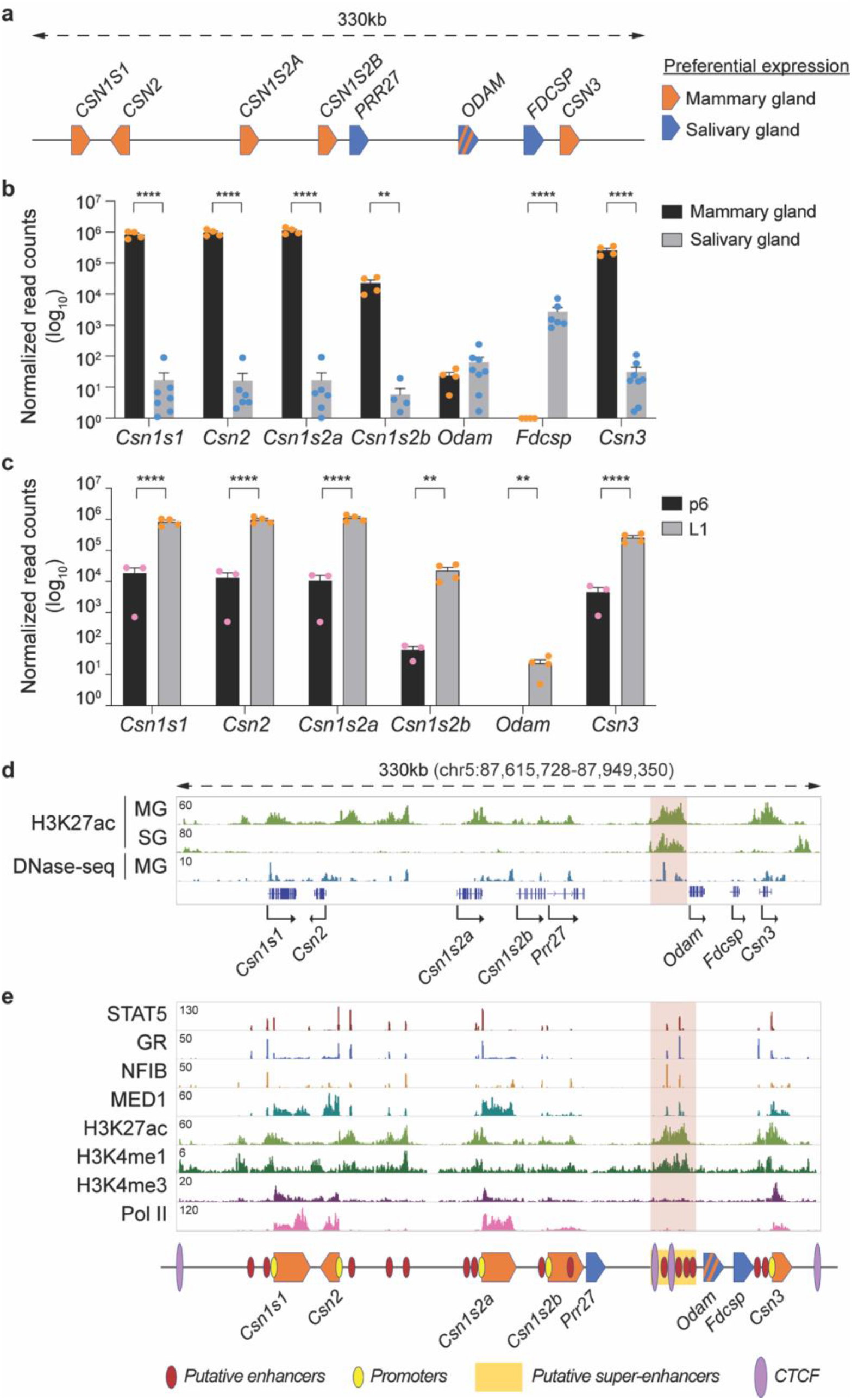
Characteristics of the *Casein* locus. **a**, Diagram presents gene structure within the *Casein* locus and their preferential expression. **b**, mRNA levels of genes in the *Casein* locus were measured by RNA-seq in lactating mammary gland and salivary gland (*n* = 4 and 8, respectively). (C) mRNA levels of *Csn* genes were measured by RNA-seq at day six of pregnancy (p6) and lactation day one (L1) (*n* = 3 and 4, respectively). **d**, Genomics feature of the *Casein* locus was identified by ChIP-seq data DNase-seq data in lactating mammary gland and salivary gland. **e**, ChIP-seq data for TFs and histone markers provided structural information on the *Casein* locus at L1. Red, yellow and purple circles are marked putative enhancers, promoters, and CTCF binding sites, respectively. Super-enhancer is indicated by yellow rectangle.

## Results

Gene activity was measured through RNA-seq conducted on lactating mammary tissue and salivary tissue (Fig. 1b). Combined, the mRNA levels of the five caseins account for more than 50% of mRNA in mammary tissue, and up to 10^6^ reads were recorded for individual *Casein* genes. *Casein* expression in salivary tissue was four orders of magnitude lower. While expression of *Fdcsp* is confined to salivary tissue, *Odam* mRNA levels are equivalent in both mammary and salivary tissue. *Casein* mRNA levels increase approximately 100-fold between day 6 of pregnancy (p6) and day 1 of lactation (L1) (Fig. 1c, Supplementary Table 1) and at least 10^4^-fold between non-parous (virgin) and day 10 lactating tissue (L10) (Supplementary Fig. 1a) (Lee et al., accompanying manuscript), making the respective genes ideal targets to understand gene activation by pregnancy hormones. The shared mammary-salivary locus also provides a unique opportunity to gain insight into control mechanisms operative in different organs. Candidate regulatory elements were identified based on H3K27ac patterns and transcription factor (TF) binding (Fig. 1d-e). As anticipated H3K27ac marks at the five *Casein* genes were restricted to mammary tissue. However, H3K27ac coverage upstream of *Odam*, the ancestral gene of the locus, was found in both mammary and salivary tissue, pointing to shared regulatory elements operative in both tissues. This region also harbors a 147 bp long evolutionarily conserved region (ECR) identified in mammals^4^.

Digging deeper, we explored binding of TFs known to activate other mammary genes^5^, such as the cytokine-inducible transcription factor Signal Transducer and Activator of Transcription (STAT5), the glucocorticoid receptor (GR), Nuclear Factor IB (NFIB) and mediator complex subunit 1 (MED1) (Fig. 1e). TF binding coincided with H3K27ac and H3K4me1 marks and a total of 20 putative regulatory elements were identified within the 330 kb locus. The H3K27ac marked region upstream of *Odam* contains four STAT5 bound regions and has all hallmarks of a super-enhancer (Fig. 1e, area highlighted in red). RNA polymerase II (Pol II) coverage was most prominent at the *Casein* genes. Integration of chromatin structures and gene expression data suggest that the mammary-salivary locus harbors the highest density of candidate enhancers and highly regulated genes among all multi-gene loci in mammary tissue (Supplementary Fig. 1b-d).

Although all 20 candidate regulatory elements were bound by STAT5, a principal TF controlling mammary development and function^6^, only 12 STAT5 peaks coincided with genuine DNA binding motifs (GAS, interferon-Gamma Activated Sequence) (Supplementary Fig. 2), suggesting STAT5 binding at other sites through alternative TFs, such as NFIB or GR (Fig. 1e). An inherent diversity in anchor proteins could lead to seemingly identical enhancers, with possibly different and unique activities. STAT5 binding occurred at the promoter regions of *Csn1s1, Csn2, Csns2a* and *Csn1s2b* and coincided with *bona fide* H3K4me1 enhancer marks^7,8^ (Supplementary Fig. 2). As expected, H3K4me3 marks were exclusively associated with promoter regions and Pol II coverage was preferentially over gene bodies (Supplementary Fig. 2).

To identify dual regulatory elements controlling genes in mammary and salivary glands, we focused on the candidate SE located between *Csn1s2b* and *Odam*, the ancestral gene expressed in both tissues. We generated mice carrying individual and combinatorial deletions of the four constituent enhancers (E) (Fig. 2a). All analyses were conducted in mammary tissue after a full pregnancy, thus monitoring hormone-induced gene activation. Deletion of the entire 10 kbp SE (ΔSE) was confirmed by the absence of TF binding and H3K27ac marks in this region (Fig. 2b). Although we had hypothesized that the entire shared locus would be under SE control, we observed distinct gene-specific differences. While *Odam* mRNA levels were reduced by more than 99%, *Csn3* by 98% and *Csn1s2b* by 93%, *Csn1s1* was reduced by a mere 50% (Fig. 2c, Supplementary Table 2) which coincided with a decline of H3K27ac, H3K4me3, and Pol II coverage at these genes (Fig. 2d, Supplementary Fig. 3). In contrast, *Csn2* and *Csn1s2a* mRNA levels were reduced at a lower, yet statistically significant, level. This experiment demonstrates that the SE dictates expression of three genes but has a limited impact on the other three mammary genes in the shared locus, whose regulation might be controlled by gene-specific enhancers.

**Fig. 2.**
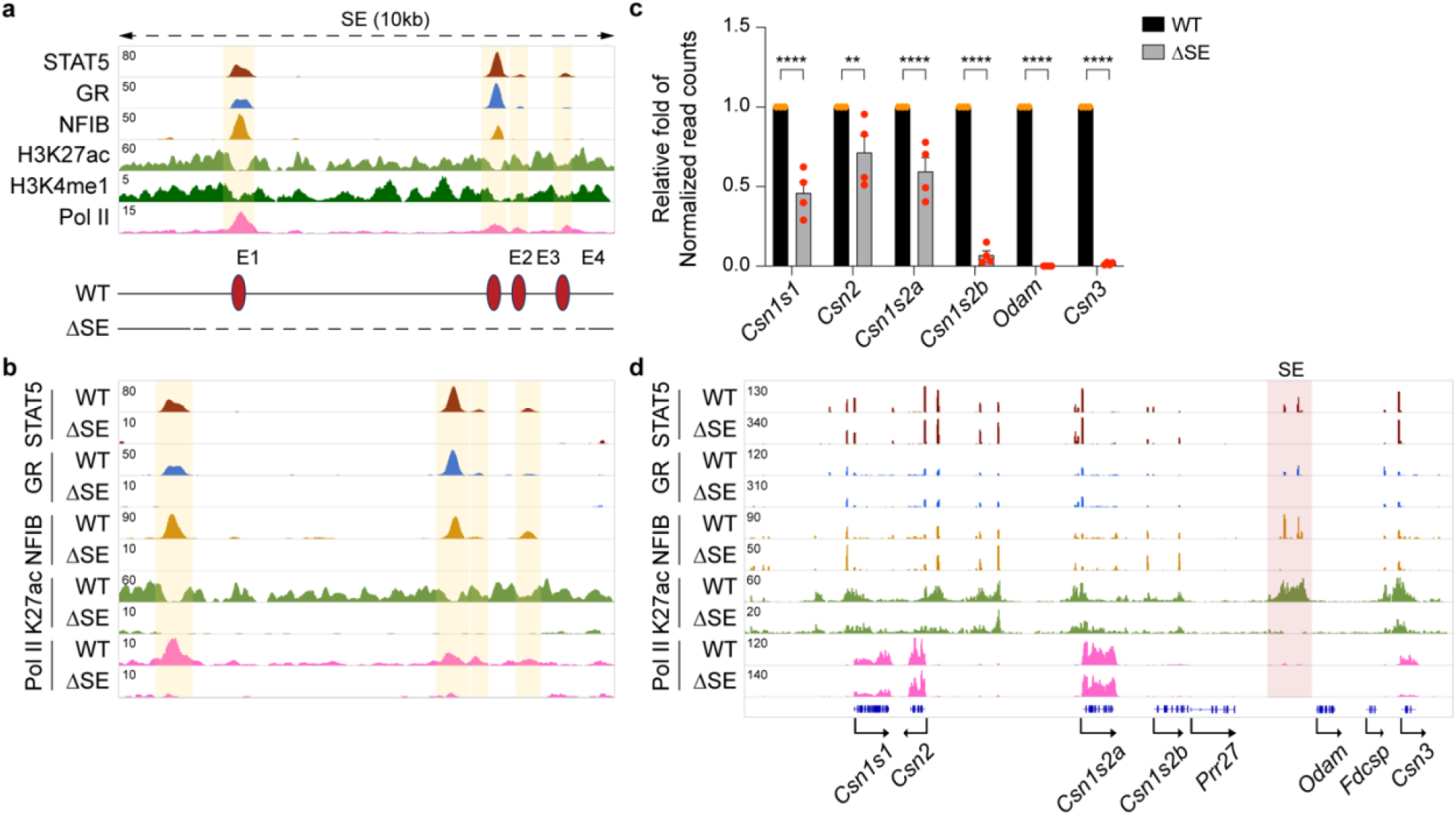
Differential activation of selected *Casein* genes by the super-enhancer during pregnancy. **a**, The putative super-enhancer was identified by ChIP-seq for TFs and activating histone marks at L1. Diagram shows the enhancer deletions introduced in mice using CRISPR/Cas9 genome editing. **b**, ChIP-seq analysis shows the genomic structure of super-enhancer in lactating mammary tissue of WT and ΔSE mice. The orange shades indicate enhancers. **c**, Expression of *Casein* genes were measured in pregnancy day 18 (p18) mammary tissue from WT and ΔSE mice by RNA-seq (*n* = 4). **d**, STAT5, GR, H3K27ac and Pol lI landscape at the *Casein* locus in WT and ΔSE tissue during pregnancy was identified by ChIP-seq.

To understand whether the SE is required for the establishment of gene-specific enhancers, we conducted additional ChIP-seq experiments. STAT5, GR and NFIB binding at the two candidate *Csn3* enhancers remained intact in the absence of the SE (Fig. 2d, Supplementary Fig. 3), suggesting the absence of compensatory activity. Notably, H3K27ac at the *Csn3* candidate enhancers depends on the SE and not the gene-specific enhancers (Fig. 2d). In contrast, STAT5 binding at the *Csn1s2b* proximal enhancer is lost in ΔSE mammary tissue suggesting that the SE activates this secondary gene-specific enhancer.

While the 10 kbp SE region harbors four individual TF peaks, their ability to function individually or in combination and contribute to the overall SE activity is not clear. To gain insight into the complexity of this SE, we introduced individual and combined deletions (Supplementary Fig. 4). Deletion of E1 (ΔE1), E2 (ΔE2) or E4 (ΔE4) resulted in the loss of STAT5 binding and H3K27ac at their respective sites. Notably, the establishment of E3 and E4 is dependent on the presence of E2. While loss of E4 (ΔE4) had no discernible consequence on any of the *Casein* genes, *Csn1s1, Csn1s2b* and *Csn3* mRNA levels were reduced between 40-70% in both ΔE1 and ΔE2 tissues. Combined deletion of both E1 and E2 (ΔE1/2) silenced the entire SE and mimicked the ΔSE mutation suggesting redundancy between E1 and E2.

A defining feature of milk protein gene is their exceptional response to pregnancy hormones, in particular prolactin. While the SE differentially affects *Casein* genes after a full pregnancy, it might have a more extended function in early pregnancy, prior to the prolactin surges that activate milk protein genes. Expression data obtained at p6 indicate an expanded SE function extending throughout the entire shared locus. In the absence of the SE (ΔSE), expression of *Odam* and all five *Casein* genes was reduced by more than 96% (Supplementary Fig. 5, Supplementary Table 3), suggesting SE affects initiation of *Casein* enhancers’ establishment and three genes that were less regulated in lactating ΔSE tissue have own enhancers established by increased hormone level during pregnancy.

Having identified the physiological significance of the SE in mammary tissue, we addressed its regulatory significance in salivary glands. While loss of SE activity led to a complete silencing of *Odam* expression, *Fdcsp* and *Csn3* mRNA levels declined by 99% and 88%, respectively (Fig. 3, Supplementary Table 4), demonstrating its dual specificity.

**Fig. 3.**
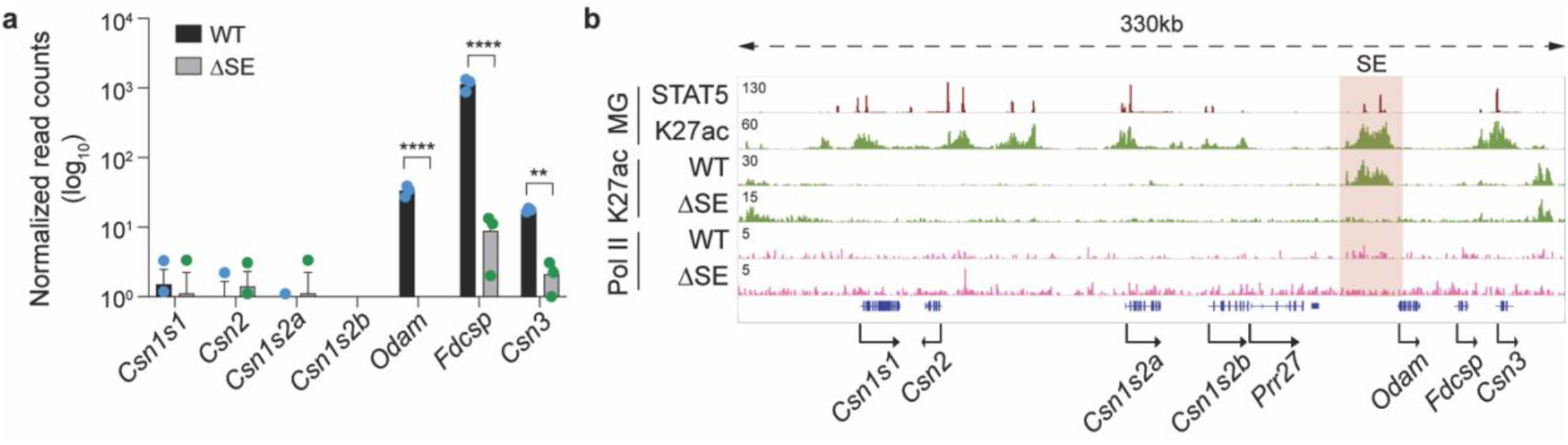
Salivary specific activation of selected genes in the *Casein* locus by the super-enhancer. **a**, Expression of *Casein, Odam* and *Fdcsp* genes was measured by RNA-seq in salivary tissue from ΔSE mice (*n* = 3). **b**, ChIP-seq analysis shows salivary landscape in WT and ΔSE mice. The red shade indicates the super-enhancer.

The absence of *Fdcsp* expression in mammary tissue might be the result of *Odam* blocking the SE from efficiently activating the *Fdcsp* promoter. To test this hypothesis, we deleted the *Odam* gene, thus transporting the SE within a few kbp to the *Fdcsp* gene (Supplementary Fig. 6, Supplementary Table 5). Despite being in the physical orbit of the SE, the *Fdcsp* remained silent in lactating mammary tissue and expression in salivary tissue was unaltered. These findings suggest that the promoter is unresponsive to the SE. In contrast, *Csn3* mRNA levels increased approximately 2-fold in mammary tissue suggesting a distance-dependency of SE activity. No expression changes were observed in salivary glands.

Removal of the SE ablates *Csn3* expression without impacting TF binding at the two *Csn3* enhancers (Fig. 2), thus questioning their physiological roles. We addressed this issue and introduced deletions within the distal (E1) and proximal (E2) candidate enhancers (Fig. 4a). Strong STAT5 binding coinciding with two GAS motifs and an NFIB site occurred at E2, and binding at E1 was weaker. GR binding was stronger at E1 compared to E2, suggesting distinct molecular structures and possibly functions of the two enhancers. While deletion of the distal candidate enhancer (ΔE1) resulted in the loss of TF binding and H3K27ac at this site, *Csn3* gene expression remained at 55% (Fig. 4b-c). In contrast, deletion of the two GAS motifs in E2 (ΔE2-S) reduced *Csn3* mRNA levels by 98%. While STAT5 binding was completely abolished, residual binding of NFIB was detected. Deletion of the two GAS motifs and the NFIB site (ΔE2-S/N) further reduced *Csn3* mRNA levels, coinciding with a complete absence of TF binding, H3K27ac and Pol II loading (Fig. 4b-c). Loss of the *Csn3* enhancer did not adversely affect other *Casein* genes (Supplementary Fig. 7, Supplementary Table 6). The combined deletion of both enhancers, E1 and E2, (ΔCsn3-E1/2) did not further reduce gene activity (data not shown) suggesting no redundancy between them. Interactions between the *Csn3* enhancers and SE had been confirmed by 3C, further supporting their crosstalk^9^.

**Fig. 4.**
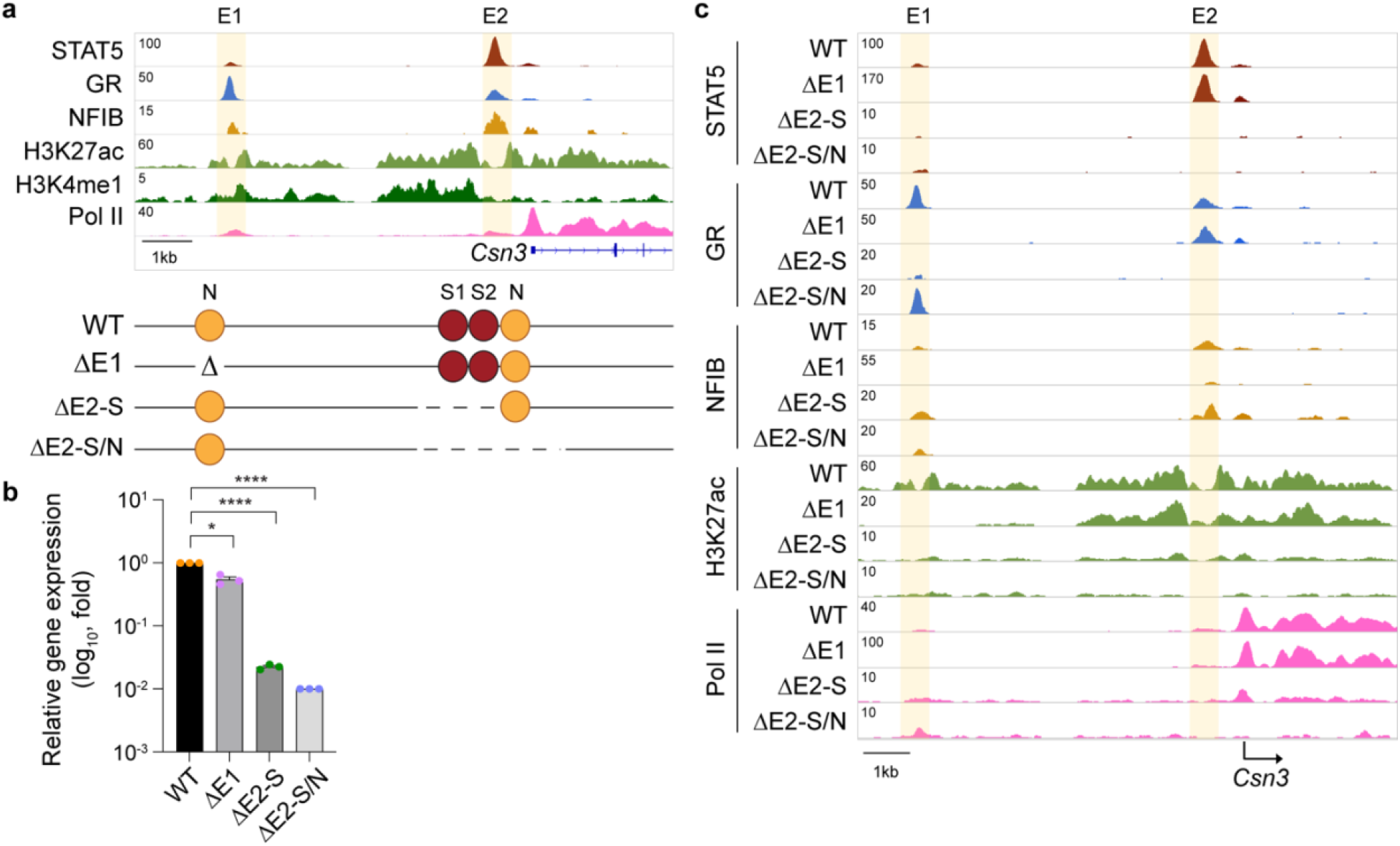
Super-enhancer-dependent gene-specific enhancers activate *Csn3* expression. **a**, the presence of H3K27ac and H3K4me1 marks indicated a distal candidate enhancer (E1) at -7 kbp and a proximal one (E2) at -0.6 kbp. Diagram shows the enhancer deletions introduced in mice using CRISPR/Cas9 genome editing. **b**, *Csn3* mRNA levels were measured by qRT-PCR in lactating mammary tissue from WT mice and mice lacking the *Csn3* distal enhancer (ΔE1) and *Csn3* proximal enhancer (ΔE2) and normalized to *Gapdh* levels. Results are shown as the means ± s.e.m. of independent biological replicates (*n* = 3). One-way ANOVA followed by Dunnett’s multiple comparisons test was used to evaluate the statistical significance of differences between WT and each mutant mouse line. **c**, Genomic features of the *Csn3* locus were investigated by ChIP-seq in lactating mammary tissue of WT, ΔE1, ΔE2-S and ΔE2-S/N mice. The highlighted orange shades indicate enhancers.

Expression of *Csn1s1*, positioned at the 5’ border of the *Casein* locus, is only marginally influenced by the distant SE, suggesting the existence of independent regulatory elements. H3K27ac marks located candidate regulatory elements at -11.5 kb (E1) and -3.5 kb (E2) (Fig. 5a). Strong STAT5, GR and NFIB occupancy was detected at site E2 but less so at E1, which also had reduced H3K27ac coverage. STAT5 binding was also observed at the *Csn1s1* promoter and coincided with a GAS motif at -100 bp. While deletion of the GR motif in E1 (ΔE1) resulted in the loss of GR and STAT5 binding at this site and reduced STAT5 binding at the promoter site, *Csn1s1* expression at lactation was unimpaired (Fig. 5b-c). In contrast, deletion of the NFIB sites in E2 (ΔE2) resulted in a 65% reduction of *Csn1s1* expression and coincided with the loss of TF binding and H3K27ac at both enhancers. These findings demonstrate that the *Csn1s1* enhancers have a very limited biological activity compared to the *Csn3* enhancer described in this study. We propose that the promoter with the STAT5 site might be the principal regulator activating *Csn1s1* expression during pregnancy.

**Fig. 5.**
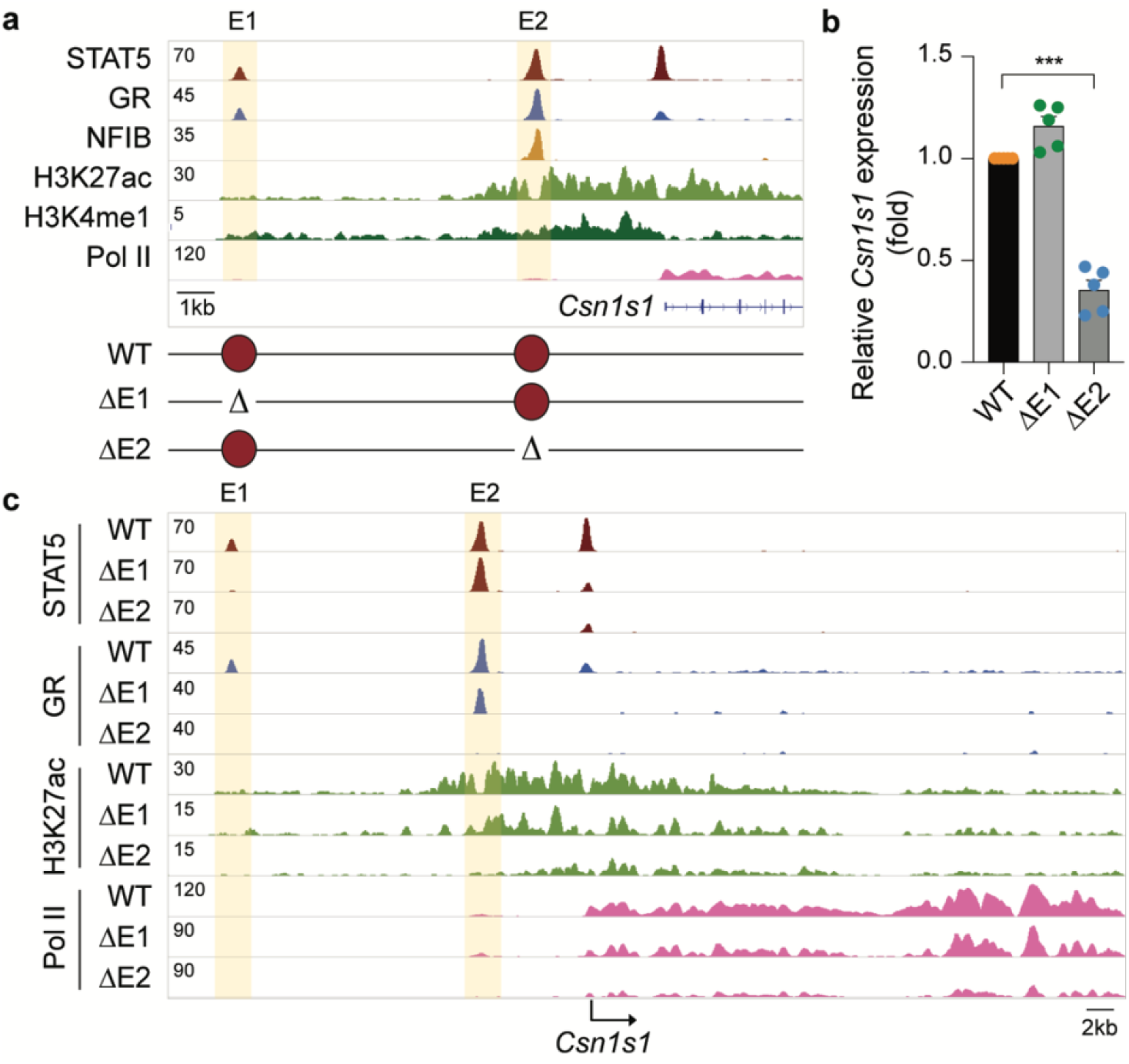
Redundant and non-redundant functions of the super-enhancer and *Csn1s1* putative enhancers in the *Csn1s1* expression. **a**, the putative Csn1s1 enhancers were identified by ChIP-seq for TFs and activating histone marks at L1. Diagram shows the deletions introduced in the mouse genome using CRISPR/Cas9 genome editing. **b**, *Csn1s1* mRNA levels in lactating mammary tissues from WT and mutant mice were measured by qRT–PCR and normalized to *Gapdh* levels. Results are shown as the means ± s.e.m. of independent biological replicates (*n* = 5). One-way ANOVA followed by Dunnett’s multiple comparisons test was used to evaluate the statistical significance of differences between WT and each mutant mouse line. **c**, The *Csn1s1* locus was profiled using ChIP-seq in WT and mutant tissue.

## Discussion

Despite a wealth of studies^5,10-12^, key questions pertaining the contribution of enhancers and SEs to gene regulation remain to be answered. Specifically, understanding the regulation of complex multi-gene loci harboring genes expressed in one or more distinct cell types is lacking. Our study provides insight into regulatory mechanisms operative in salivary and mammary glands, tissues that share morphological and molecular features during embryogenesis^13-15^. Specifically, we identified a SE exclusively active in mammary and salivary tissue.

The shared locus with its eight genes linked to lactation, saliva and immune response^1^ is an evolutionary playground that fostered regulatory innovation and yielded 20 enhancer and promoter elements. *Odam* and its associated SE likely constitute the ancestral unit of the shared locus and regulatory activity expanded from salivary tissue to mammary tissue. However, as this locus expanded, the five newly formed *Casein* genes acquired their own regulatory elements and three gained independences of the SE. Although *Csn1s2b*^*9*^ and *Csn3* acquired distal enhancers essential for their expression in lactating mammary tissue, they still retained their dependence on the SE. The enhancers linked to the five *Casein* genes display equivalent structures and TF occupancies suggesting the presence of additional elements that facilitate SE independence of three *Casein* genes. ChIP-seq analyses identified the presence of cytokine-response elements within the promoter regions of these three *Casein* genes and mutational analyses determined these as critical elements in at least the *Csn2* gene (Lee et al., accompanying manuscript). Unlike the shared mammary-salivary locus, expression of the five globin genes in the α-globin locus is dependent on one SE composed of five individual erythroid enhancers^12,16^. Moreover, the two SE exhibit mechanistic differences, with enhancer redundancy in the mammary-salivary locus and additive activity in the α-globin locus.

While genetic studies have identified key transcription factors controlling mammary-specific gene expression through enhancers and SEs, there is limited knowledge about regulatory mechanisms operative in salivary tissue. Genome-wide histone modification studies in mouse submandibular glands have pointed to putative regulatory regions^17^, but no defined salivary-specific enhancers have been described. Key TFs controlling mammary function, such as STAT5^6,18^ and ELF5^19^ are dispensable in the salivary gland^20^. At this point the molecular backbone of the salivary enhancer, as defined by H3K27ac marks, extends over 10 kbp but no associated salivary-specific TF have been identified. Also, SE-induced gene activation in salivary tissue is lower than in mammary tissue.

The *Fdcsp* gene is silent in mammary tissue and uniquely activated in salivary glands by the SE suggesting a cell-preferential response of its promoter. This specificity is not influenced by distance of the SE or the presence of the intervening *Odam* gene, suggesting the presence of unique promoter elements that permit enhancer sensing. Alternatively, differential promoter accessibility^21^ in salivary and mammary tissue could account for the cell specificity.

Here we report on regulatory innovation in an evolutionary playground with genes acquiring mammary and salivary specificity (Fig. 6). The elaborate enhancer structure developed in the mammary lineage permits an exceptional expression of five genes accounting for 80% of milk proteins, an essential requirement for the sustained success of mammals. We propose that the concentration of enhancers and their high-density occupation with TF and co-activators provides an optimal regulatory environment. The co-existence and interdependence of a SE and gene-specific enhancers provides opportunities for the *Casein* locus to rapidly develop and produce milks with vastly different properties.

**Fig. 6.**
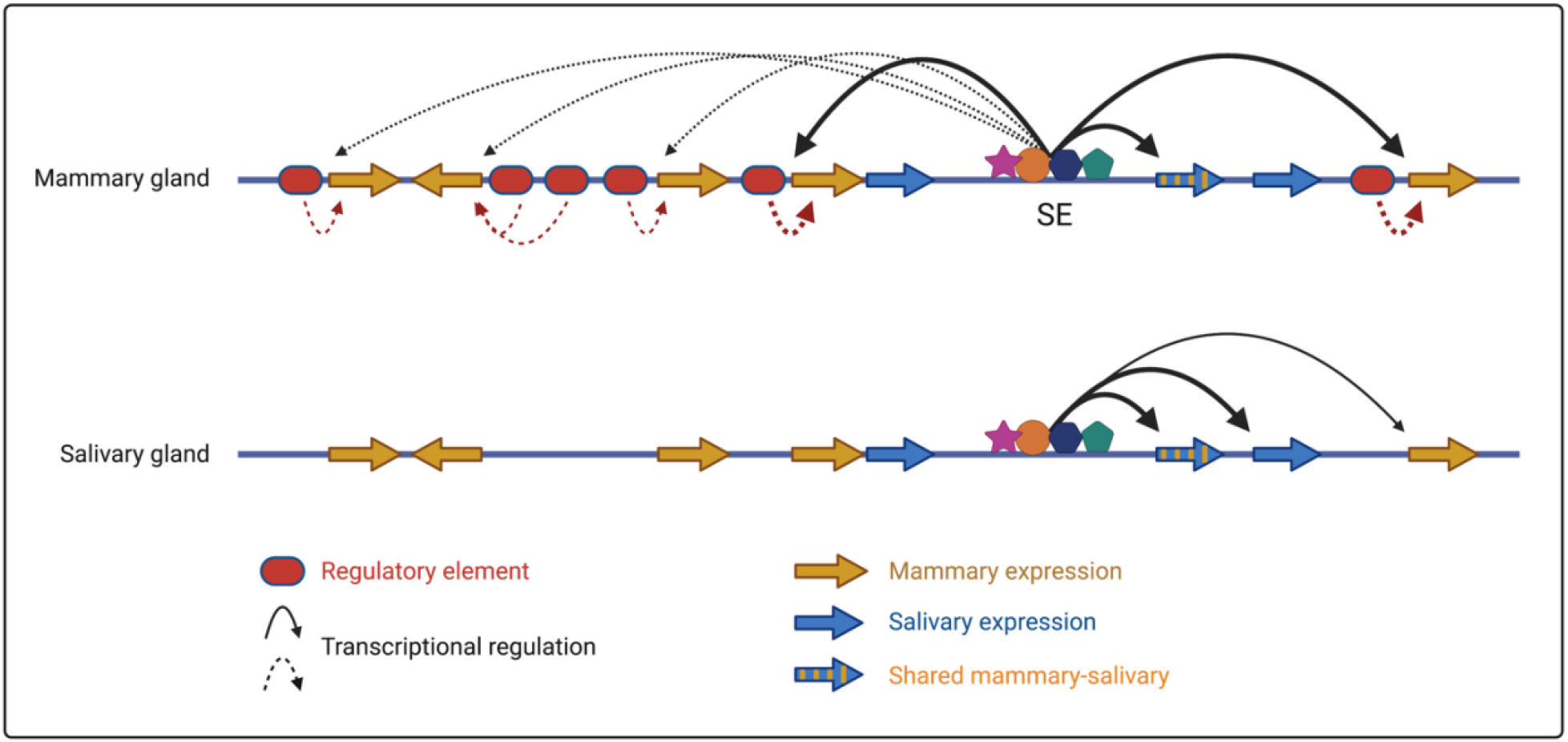
Proposed model outlining regulation of the shared mammary-salivary by a dual specific super-enhancer and gene-specific enhancers. Super-enhancer preferentially activates the *Csn1s2b, Csn3* and *Odam* genes and marginally the *Csn1s1, Csn2* and *Csn1s2a* genes during pregnancy in mammary gland and regulates the promoters of *Odam, Fdcsp* and *Csn3* genes in salivary gland.

## Materials and Methods

### Mice

All animals were housed and handled according to the Guide for the Care and Use of Laboratory Animals (8th edition) and all animal experiments were approved by the Animal Care and Use Committee (ACUC) of National Institute of Diabetes and Digestive and Kidney Diseases (NIDDK, MD) and performed under the NIDDK animal protocol K089-LGP-17. CRISPR/Cas9 targeted mice were generated using C57BL/6N mice (Charles River) by the transgenic core of the National Heart, Lung, and Blood Institute (NHLBI). Single-guide RNAs (sgRNA) were obtained from either OriGene (Rockville, MD) or Thermo Fisher Scientific (Supplementary Table 7). Target-specific sgRNAs and *in vitro* transcribed *Cas9* mRNA were co-microinjected into the cytoplasm of fertilized eggs for founder mouse production. The ΔE1/2 mutant mouse was generated by injecting a sgRNA for E2 into zygotes collected from ΔE1 mutant mice. All mice were genotyped by PCR amplification and Sanger sequencing (Macrogen and Quintara Biosciences) with genomic DNA from mouse tails (Supplementary Table 8).

### Chromatin immunoprecipitation sequencing (ChIP-seq) and data analysis

Mammary tissues from specific stages during pregnancy and lactation were harvested, and stored at -80°C. The frozen-stored tissues were ground into powder in liquid nitrogen. Chromatin was fixed with formaldehyde (1% final concentration) for 15 min at room temperature, and then quenched with glycine (0.125 M final concentration). Samples were processed as previously described^22^. The following antibodies were used for ChIP-seq: STAT5A (Santa Cruz Biotechnology, sc-1081 and sc-271542), GR (Thermo Fisher Scientific, PA1-511A), MED1 (Bethyl Laboratory, A300-793A), H3K27ac (Abcam, ab4729), RNA polymerase II (Abcam, ab5408), H3K4me1 (Active Motif, 39297) and H3K4me3 (Millipore, 07-473). Libraries for next-generation sequencing were prepared and sequenced with a HiSeq 2500 or 3000 instrument (Illumina). Quality filtering and alignment of the raw reads was done using Trimmomatic^23^ (version 0.36) and Bowtie^24^ (version 1.1.2), with the parameter ‘-m 1’ to keep only uniquely mapped reads, using the reference genome mm10. Picard tools (Broad Institute. Picard, http://broadinstitute.github.io/picard/. 2016) was used to remove duplicates and subsequently, Homer^25^ (version 4.8.2) and deepTools^26^ (version 3.1.3) software was applied to generate bedGraph files, seperately. Integrative Genomics Viewer^27^ (version 2.3.81) was used for visualization. Coverage plots were generated using Homer^25^ software with the bedGraph from deepTools as input. R and the packages dplyr (https://CRAN.R-project.org/package=dplyr) and ggplot2^28^ were used for visualization. Each ChIP-seq experiment was conducted for two replicates. Sequence read numbers were calculated using Samtools^29^ software with sorted bam files. The correlation between the ChIP-seq replicates was computed using deepTools using Spearman correlation.

### Total RNA sequencing (Total RNA-seq) and data analysis

Total RNA was extracted from frozen mammary tissue from wild-type mice at day six of pregnancy and purified with RNeasy Plus Mini Kit (Qiagen, 74134). Ribosomal RNA was removed from 1 μg of total RNAs and cDNA was synthesized using SuperScript III (Invitrogen). Libraries for sequencing were prepared according to the manufacturer’s instructions with TruSeq Stranded Total RNA Library Prep Kit with Ribo-Zero Gold (Illumina, RS-122-2301) and paired-end sequencing was done with a HiSeq 2500 instrument (Illumina).

Total RNA-seq read quality control was done using Trimmomatic^23^ (version 0.36) and STAR RNA-seq^30^ (version STAR 2.5.3a) using paired-end mode was used to align the reads (mm10). HTSeq^31^ was to retrieve the raw counts and subsequently, R (https://www.R-project.org/), Bioconductor^32^ and DESeq2^28^ were used. Additionally, the RUVSeq^33^ package was applied to remove confounding factors. The data were pre-filtered keeping only those genes, which have at least ten reads in total. Genes were categorized as significantly differentially expressed with an adjusted p-value below 0.05 and a fold change > 2 for up-regulated genes and a fold change of < -2 for down-regulated ones. The visualization was done using dplyr (https://CRAN.R-project.org/package=dplyr) and ggplot2^34^.

### RNA isolation and quantitative real-time PCR (qRT–PCR)

Total RNA was extracted from frozen mammary tissue of wild type and mutant mice using a homogenizer and the PureLink RNA Mini kit according to the manufacturer’s instructions (Thermo Fisher Scientific). Total RNA (1 μg) was reverse transcribed for 50 min at 50°C using 50 μM oligo dT and 2 μl of SuperScript III (Thermo Fisher Scientific) in a 20 μl reaction. Quantitative real-time PCR (qRT-PCR) was performed using TaqMan probes (*Csn1s1*, Mm01160593_m1; *Csn2*, Mm04207885_m1; *Csn1s2a*, Mm00839343_m1; *Csn1s2b*, Mm00839674_m1; *Odam*, Mm02581573_m1; *Csn3*, Mm02581554_m1; mouse *Gapdh*, Mm99999915_g1, Thermo Fisher scientific) on the CFX384 Real-Time PCR Detection System (Bio-Rad) according to the manufacturer’s instructions. PCR conditions were 95°C for 30s, 95°C for 15s, and 60°C for 30s for 40 cycles. All reactions were done in triplicate and normalized to the housekeeping gene *Gapdh*. Relative differences in PCR results were calculated using the comparative cycle threshold (*CT*) method and normalized to *Gapdh* levels.

### Identification of regulatory elements

MACS2^35^ peak finding algorithm was used to identify regions of ChIP-seq enrichment over the background to get regulatory elements at L1 and L10. Peak calling was done on both STAT5A replicates and broad peak calling on H3K27ac. Only those peaks were used, which were identified in both replicates and with H3K27ac coverage underneath.

### Identification of complex mammary loci

Mammary specific genes were identified using RNA-seq data from pregnancy day six (p6), lactation day one (L1) and ten (L10). Those genes were considered, which were induced more than two-fold with an adjusted p-value below 0.05 between p6 and L1 or p6 and L10. The next step comprised the stitching of neighboring genes, by only considering protein coding genes. Those stitched loci were subsequently compared to the contact domains (Hi-C data) and only those loci passed the validation that were not overlapping with any border of the contact domains. If a locus overlapped they were treated the following way: (i) if the locus contains only two genes it was discarded, as it will not pass the prerequisite that a complex locus comprises at least two genes; (ii) all other loci were split up at the border and only those were kept that comprised more than two genes; (iii) the loci were shrunk to the size of the remaining genes. As possible regulatory elements (in our analysis STAT5A) are also part of complex loci, we expanded the borders of each locus to comprise STAT5A binding sites, if they were located within the adjacent intergenic region. Those new loci were finally checked again to not overlap with contact domain boundaries, otherwise they were shrunk down to the last element not overlapping with the contact domain; (iv) the final list of complex loci comprises loci with at least three genes.

The analysis was done using bedtools^36^, bedops, R (https://www.R-project.org/) and Bioconductor^32^ as well as the R packages dyplr (https://CRAN.R-project.org/package=dplyr) and ggplot2^37^.

### Statistical analyses

All samples that were used for qRT–PCR and RNA-seq were randomly selected, and blinding was not applied. For comparison of samples, data were presented as standard deviation in each group and were evaluated with a *t-*test and 2-way ANOVA multiple comparisons using PRISM GraphPad. Statistical significance was obtained by comparing the measures from wild-type or control group, and each mutant group. A value of \**P* < 0.05, \*\**P* < 0.001, \*\*\**P* < 0.0001, \*\*\*\**P* < 0.00001 was considered statistically significant. ns, no significant.

## Data availability

All data were obtained or uploaded to Gene Expression Omnibus (GEO). ChIP-seq data of wild-type tissue at L1 and L10 were obtained under GSE74826, GSE119657 and GSE115370. RNA-seq data for WT at L1 as well as L10 were downloaded from GSE115370. The RNA-seq data for WT and mutant mice at p6 and p18, Hi-C and 4C-seq data for WT and mutant mice at L1 and ChIP-seq data for WT and mutant mice were uploaded to GSE127144 (ChIP-seq in GSE127139, RNA-seq in GSE127140). Reviewer link will be shared upon request.

## Acknowledgments

We thank Ilhan Akan, Sijung Yun and Harold Smith from the NIDDK genomics core for NGS. This work utilized the computational resources of the NIH HPC Biowulf cluster (http://hpc.nih.gov). This work was supported by the Intramural Research Programs (IRPs) of National Institute of Diabetes and Digestive and Kidney Diseases (NIDDK) and National Heart, Lung, and Blood Institute (NHLBI).

## Author contributions

H.K.L. and L.H. designed the study and analyzed data. C.L. generated mutant mice. H.K.L. established mutant mouse line. H.K.L. performed experiments and data analysis. M.W. identified multi-gene loci. H.K.L. and L.H. supervised the study. H.K.L. and L.H. wrote the manuscript and all authors approved the final version.

